# Carbon Nanotube Electrodes for Electrochemical Detection of Dopamine

**DOI:** 10.1101/2021.08.24.457511

**Authors:** Gaurang Khot, Frank Platte, Neil Shirtcliffe, Tansu Celikel

## Abstract

Carbon nanotubes (CNTs) are suited for neurochemistry because of their biological inertness, ability to withstand biofouling, and superior electron transport kinetics. Dopamine, the canonical monoaminergic neuromodulator, contributes to reward, cognition and attention, however, its detection in real-time is challenging due to its low basal concentration in the brain (100nM L^-1^). In our present work, we fabricate pyrolytic carbon electrodes and perform a CNT coating to improve the electrochemical kinetics of dopamine. Upon CNTs coating, dopamine shows a sensitivity of 9±18nA/*μ*M for a cylindrical electrode having a mean surface diameter of 8±4μm. Increasing the scan frequency from 10-100 Hz shows that dopamine electron transfer kinetics improves; wherein dopamine is oxidized at 0.35±0.09V and reduced to -0.10±0.05V for 10 Hz. Increasing the frequency results in a shift of oxidation peak towards the anodic region, wherein dopamine oxidizes at 0.08±3V and reduces at -0.1±0.05V for 100 Hz, thus showing that dopamine redox is reversible which can be attributed to the superior electron transport kinetics of CNTs. The sensor was able to distinguish dopamine signals against other neurochemicals like serotonin and foulant 3,4-Dihydroxyphenylacetic acid (DOPAC). The minimum chemical detection that can be performed using these nanopipettes is 50±18nM L^-1^, which is well below the physiological concentrations of dopamine in the brain.

**Graphical Abstract:** **A:** Pictorial view of background-subtracted voltammetry. The waveform used was -0.4V to 1.3 V and cycled back to -0.4V at 10 Hz. **B:** The voltammogram was converted as a 2-D representation, into current, voltage, and repetition to understand the dopamine oxidation. **C:** Background subtracted voltammetry for dopamine using 100 Hz waveform. **D:** The 2-D representation of current, voltage, and repetition.

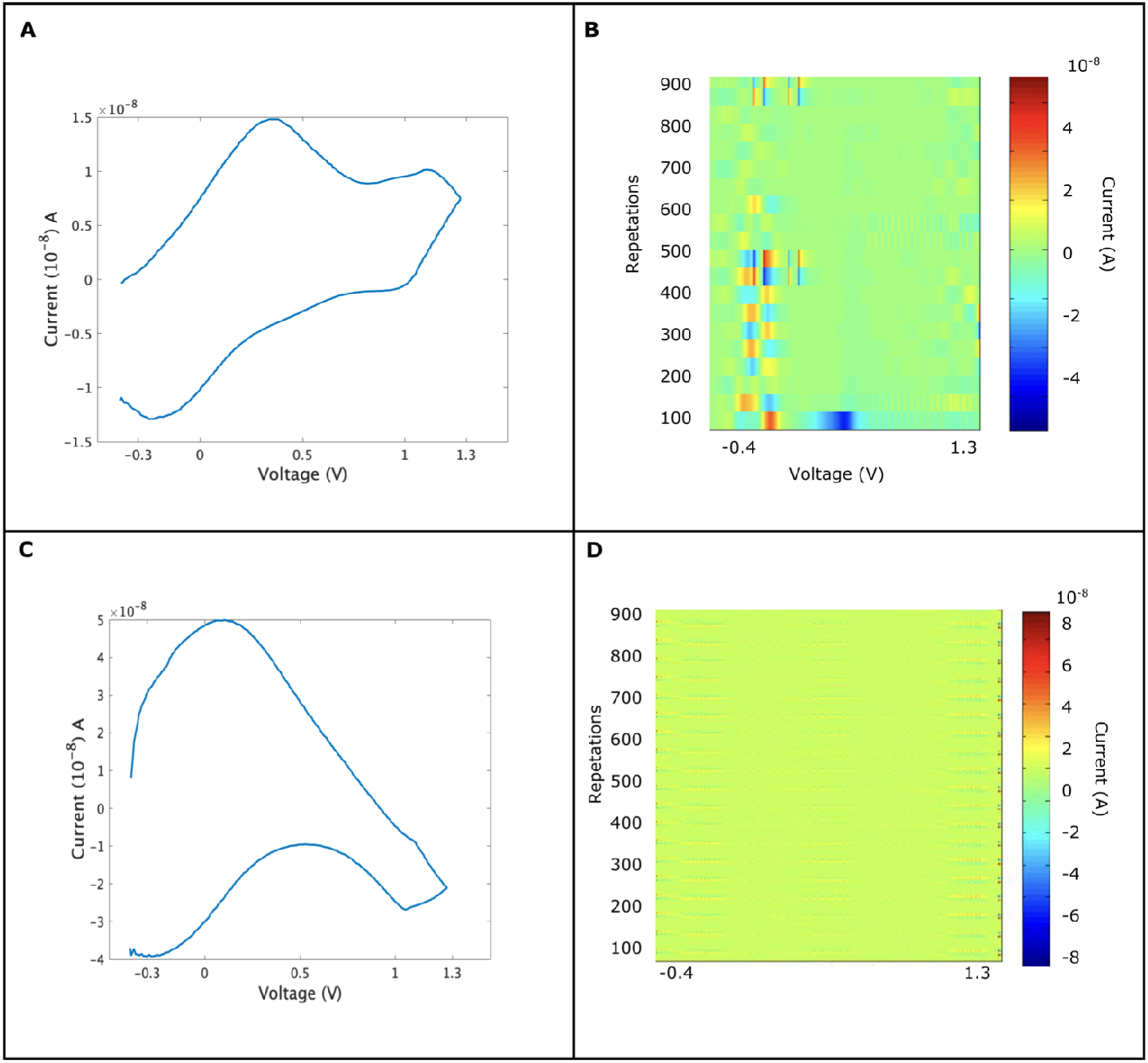

## Introduction

Dopamine is a monoaminergic neuromodulator that is responsible for reward, cognition, and motivation^1–4^. Low levels of dopamine in the brain are responsible for cognitive, motor, and behavioral disorders such as Alzheimer’s, Parkinson and Huntington’s diseases^1,3^. Dopamine regulation in the brain occurs in a sub-second time frame^5^, thus requiring sensors that have superior temporal responses and are able to measure dopamine in sub-second time frames^6^. Electrochemical sensors are widely used *in vivo* due to their ability to measure the target analyte in sub-second time frames^5,7^ The application of fast scan rate with carbon fiber microelectrodes which can fit into tight biological matrices has allowed detection of neurotransmitters and neuromodulators^6,8^. The drawback of a carbon fiber probe is biofouling and degradation of the probe over time thus limiting its application as a long-term implant material. Hence a need exists to develop materials that can serve as long-term implants in the brain.

Carbon nanotubes (CNTs) have received considerable attention in biology because of their miniature size, bio inertness, and ability to withstand befouling^9^. Their long-term bio-stability and their ability to provide faster electron kinetics have received considerable attention in electronics, semiconductors, and biochemical sensor development^9–11^. Their ability to form long sp^2^ bonds of graphite sheets in supercoiled 3--dimensional long chains, giving them superior mechanical strength than other composite materials^9,12,13^. These unique properties of CNTs have attracted the attention of neuroscientists who wish to employ CNT-based materials as neurological implants.

Fast Scan Cyclic Voltammetry (FSCV) allows measurements of compounds on a sub-second time-frame^14,15^ thus the method is ideal for molecules like neurotransmitters that have a short half life. CNTs based microelectrodes such as carbon-nanotube yarn^10,16–18^ or chemical modification of polymers, e.g. polymers blending with carbon nanotubes, have been used for electrochemical detection of dopamine^19–22^ given their ability to resist biofouling and perform repetitive measurements at faster scan rates. Advantages of CNTs materials are reduced potentials^10^ and structural stiffness^19^ making them a suitable candidate for implantation in harsh biological matrices like the brain. While carbon fiber probes are considered as a gold standard material for in vivo and in vitro electrochemistry^15^, the slow electrochemical kinetics in neurotransmitter detection limits the temporal resolution of the signals acquired using carbon fiber electrodes.

Considerable progress has been made towards electrochemical detection of dopamine on CNTs electrodes^10^, however little is understood about the oxidation kinetics of dopamine on CNTs electrodes. Thus a need exists to develop CNTs based material that can allow detection of dopamine with improved electrochemical performances. Herein we employed a CNT coating onto pyrolytic electrodes for electrochemical detection of dopamine. We investigate whether CNT coatings improves sensitivity and the rate of electron transfer for dopamine oxidation. Using our experimental data, we investigate the heterogeneous redox nature of dopamine onto pyrolytic coated CNT electrodes. Finally, we construct a Density Function Theory (DFT) model to visualize the binding of dopamine onto the CNT surfaces.

## Results and Discussions

In our earlier work, we found experimental evidence of 2 electron oxidation mechanisms for dopamine and our DFT studies showed that heterogeneous graphite surfaces show the possibility of electron and proton transfer (Khot et al., in submission). While Wightman and co-workers provide a possible mechanism for the oxidation of dopamine by 2 electrons and 2 proton transfer^7,14^. Compton’s group provided a theoretical mechanism for electron-proton transfer model for electrochemical oxidation of dopamine on gold and carbon electrodes at acidic and neutral pH values^23,24^. In this work, we investigate if we can make use of CNT coatings on pyrolytic electrodes, which should increase the sensitivity and reduce the potential required for dopamine detection.

### 1. Electrochemical Characterization of Pyrolyzed Electrode Coated with CNTs for FSCV

To investigate the electrochemical properties of the CNT coated electrodes (**Figure 1A**), a flow cell was set up that mimics the transient change of dopamine in the brain. Probes were pyrolyzed for 60 s, to obtain sub-micrometer surface area and dip-coated with CNTs suspended in Nafion®. Background current was measured and if it was higher than 1.1μA, the electrodes were discarded.

**Figure 1:**
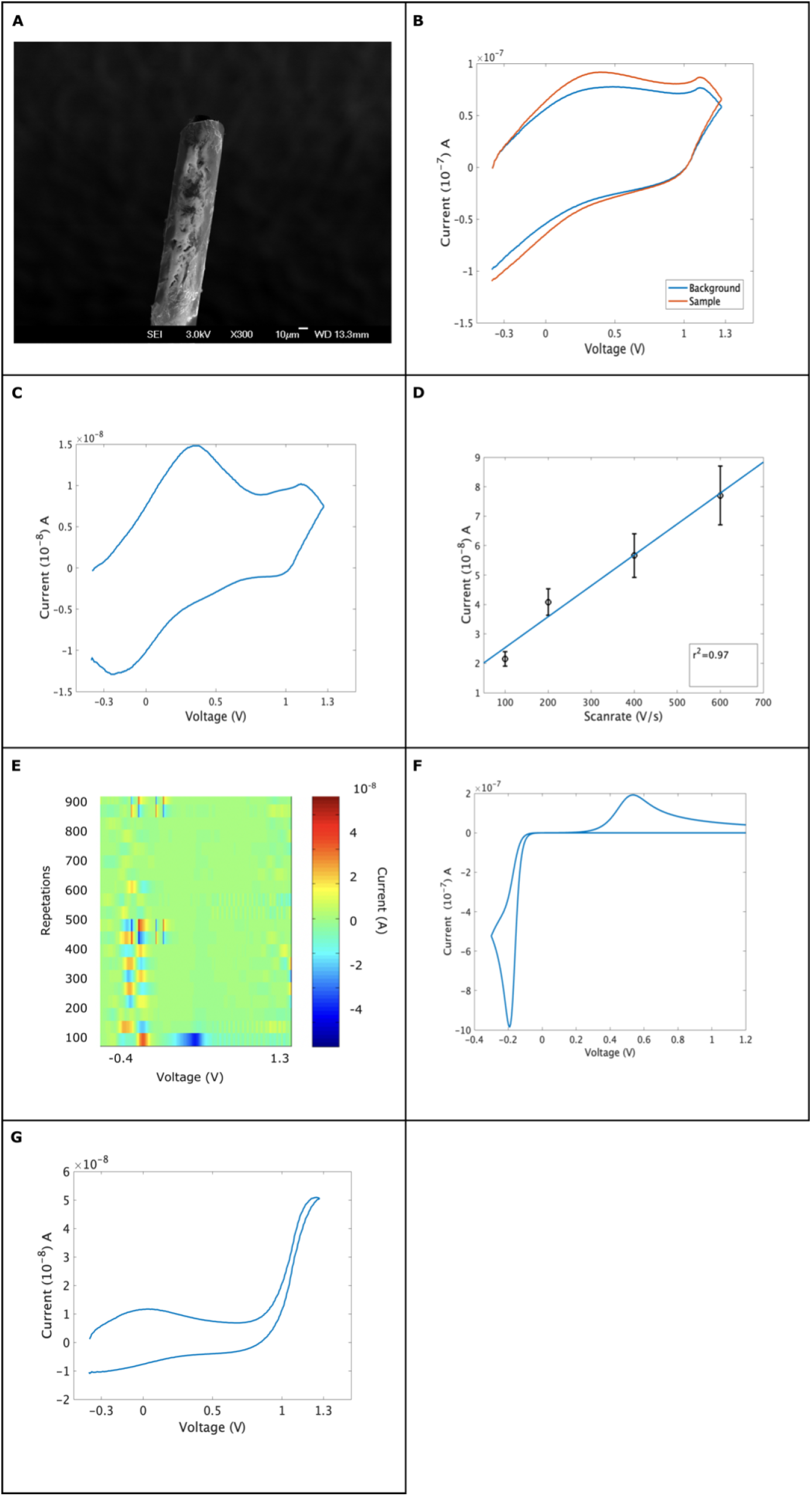
The characterization of dopamine on CNT probes. **A**: Shows the micrograph of pyrolytic electrode heated for 60 seconds and coated with CNTs suspended in the Nafion membrane. The coating had a mean diameter of 5μm and a nominal height of 0.9μm. **B**: Shows the overlaying of the background buffer and the sample. To get the true Faradaic signal of the molecule, digital background subtraction is done. The difference in background current and the background-subtracted voltammogram is not significant (unpaired t-test, where P value 0.356, for p<0.01). **C**: Shows the result of background subtraction for 1*μ*M of dopamine, where the waveform used was -0.4V to 1.3V and cycled back to -0.4V at 10 Hz. Dopamine is oxidized at 0.35±0.09V and reduced at -0.15±0.05V. **D**: Shows scan rate against peak current, the r^2^ =0.97 showing adsorption as a mode of transport. **E**: Shows current across trials wherein the yellow font shows the oxidation of dopamine around 0.35V and a blue font shows a reduction of dopamine. A secondary reaction can be seen at vortex potential. The current generated for a circular electrode that has a mean diameter of 5μmeter is 1.2nanomole/micrometer. **F:** Shows the simulated cyclic voltammogram for a disk-shaped electrode, having a mean diameter of 5μmeter. The parameters used for 1in the simulations is diffusion constant for dopamine (D)= 0.76 × 10^--6^ cm^2^ s^-1^using electron rate transfer coefficient as 3X 10^-6^ cm^2^ s^-1^ and number of electrons transported is (n)=2. The parameters described are as described before^10^. **G**: Shows 1μm of dopamine with 200 μm of hydrogen peroxide, which is sampled using the parameters identical to that of dopamine detection (It should be noted that the anodic vortex limit was 1.3V).

To investigate dopamine dynamics, “Dopamine waveform”^25^ was applied to pyrolyzed-CNT electrodes, from -0.4V to 1.3V and cycled back to -0.4V at a repetition rate of 10Hz and the scan speed of 400V s^-1^ with a sampling frequency of 50kHz. (**Figure 1B**) shows the background current overlaid onto the sample for 1*μ*M of dopamine. The background-subtracted voltammogram for 1*μ*M of dopamine (**Figure 1C**) shows that dopamine is oxidized at 0.35V and reduces at -0.2V In addition to dopamine oxidation, secondary oxidation is seen near the anodic vertex potential at 1.1V, which we postulate is the deprotonation of dopamine caused by electrochemical oxidation of hydroxyl functional groups, located on the catechol ring. We hypothesize that the intermediate radicals reacts with the aqueous environment to form a peroxide-like species which reduces the catheol group at 0.9V Surprisingly the kinetics for secondary redox peaks were found to be reversible, wherein the difference of oxidation to reduction peak was less than 57mV, while such property was not seen for dopamine. While such observations of secondary peaks have been reported for oxidation of aminopurines^26^

The occurrence of secondary redox peaks for dopamine can be seen on carbon nanotube yarn surfaces that are laser-treated^16^. It should be noted that such secondary oxidation peaks are only seen on nanotube surfaces that have undergone a secondary physical treatment to improve their sensitivity, rate of electron transport process^17,18^. In our work, we did not perform any secondary treatment to our surface or CNTs, yet we have improved the rate of electron transfer (**Figure 1F**). Studies have shown co--detection of dopamine and hydrogen peroxide on carbon fiber electrodes, wherein dopamine oxidizes at +0.6V and reduces at -0.4V^27^, while hydrogen peroxide oxidizes at 1.3V^28^. As we intended to detect dopamine alone we did not employ a catalase enzyme-based assay to observe the decomposition of the peak at 1.1V. However, based on the peak location and the rise of signal amplitude when hydrogen peroxide is added to the solution (**Figure 1G**) it remains likely that secondary peak can be hydrogen peroxide.

To study the transport kinetics of dopamine, the scan rate was varied keeping the concentration of dopamine constant 1*μ*M. Oxidation peak current (iP) was plotted as a function of scan rate (V) and a function of the square root of scan rate (V^1/2^) to understand the transport kinetics^18^. At concentrations of 1*μ*M, scan rate and fits, with peak current, thus suggesting that dopamine is predominantly adsorbed on the surface of the electrodes (**Figure 1D**). However, once the sites are occupied, subsequent interactions of dopamine from the bulk to the surface of the electrodes occur via the diffusion of dopamine on the surface of the electrode.

### 2. Manipulating Sampling frequency

In our next set of experiments, we investigated whether alterations in sampling frequency improves the rate of electron transport for dopamine. We employed different scan frequencies; 10 Hz, 20 Hz, 40 Hz, 80 Hz, and 100 Hz and kept the scan rate constant at 400V s^-1^. The oxidation of dopamine at 400Vs^-1^ 10Hz (**Figure 2A**) and T-400V s^-1^ 20 Hz (**Figure 2B**) shows no reversible property, however a change in amplitude for secondary oxidation peak is observed, wherein the ratio of primary to secondary signal, the amplitude of secondary peak increases. A possible reason can be, as the scan frequency increases the rate of dopamine oxidation along with a non-linear rate of deprotonation increase. As the scan frequency was increased to 40 Hz, a shift towards the anodic potential for dopamine was observed, wherein dopamine oxidized at 0.18V, the reduction peak remained constant (**Figure 2C**). The amplitude of the secondary peak is increased. It must be noted that the reaction kinetics for dopamine are starting to be reversible, i.e the ratio of oxidation to reduction peak is approximately 57mV. As the scan frequency was increased to 80 Hz, dopamine oxidized to 0.08V and reduced to -0.05V, thus making the overall reaction reversible (**Figure 2D**). A similar result was seen for 100 Hz, where dopamine is oxidized at similar potentials, however, a large increase in peak current was observed (**Figure 2E**). We did not measure an increase in amplitude for the peroxide peak at 1.1V beyond 40Hz (**Figure 2C--2E**), thus suggesting that the formation of secondary species is limited by oxidation rate, and the time course for oxidation of dopamine and generation of hydrogen ions, is a time-dependent rate-limiting step.

**Figure 2:**
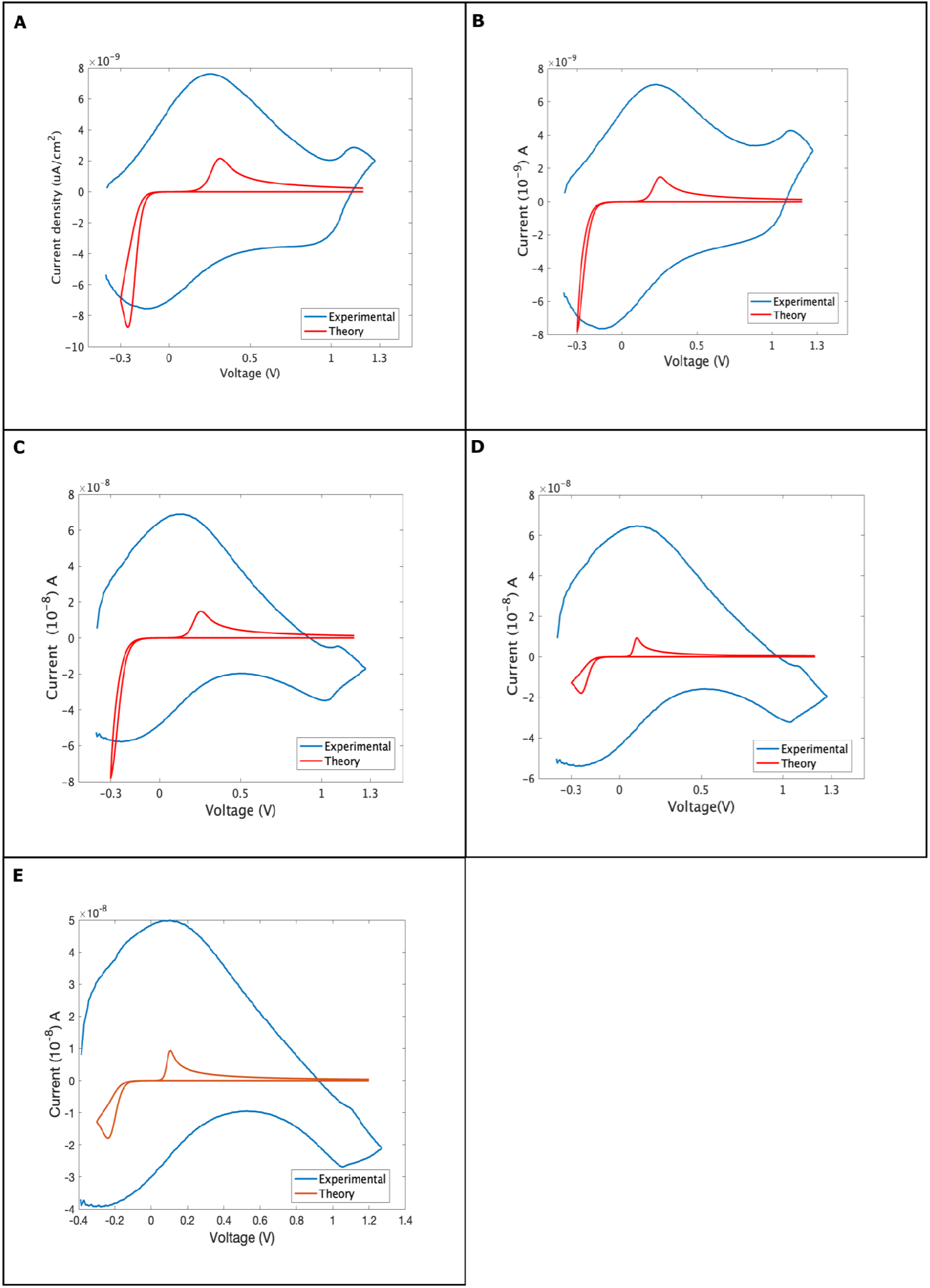
Cyclic voltammograms of 1*μ*M of dopamine sampled at different frequencies (10-100Hz), at 400V s^-1^. The experimental fit had constant D for dopamine was taken as 7.6 × 10^-6^ cm^2^ s^-1^, the area of the electrode was 0.0001 cm, α=1. The number of electrons transferred was 4, the rate of electron transfer (k) was 0.000003cm/s, the scan speed was 200 V s^-1^ and the temperature was 310 K. **A**: Shows the oxidation of dopamine at 0.35V and reduction at -0.15V. **B:** Shows the oxidation of dopamine at 0.3V while no change in reduction properties is observed. **C:** Shows a shift of dopamine oxidation on cathodic potential, wherein dopamine oxidize at 0.28V and is reduced at -0.17V. **D**: Shows the oxidation of dopamine at -0.15V and reduction at -0.2 V., thus showing reversible kinetics for dopamine. **E**: The oxidation potential of dopamine is at 0.1V and reduction is at -0.2V thus showing reversible kinetics for dopamine.

Using our data from scan frequency, we updated the theoretical modelling data of “Electron-Proton” given by Compton’s group^23,24^. As the scan frequency is increased the current increases as well, and the reaction becomes reversible. For scan frequencies of 80Hz and 100Hz, the reaction is reversible while it remains nearly reversible for 40 Hz. The rate of electron transfer depended on the scan frequency, increasing from 4 × 10^-4^ cm^2^ s^-110^ to 3 × 10^-3^ cm^2^ s^-1^ for scan frequencies of 80Hz and 100Hz. Although we did not measure a change in the rate of the electron transfer when only the scan rate alone was increased. Thus showing that the reversibility of the dopamine signature is dependent on scan frequency and not scan rate.

Previous studies using different scan frequencies onto CNT surfaces report the reduced potential required for dopamine detection, however they did not report the reversibility of dopamine electrochemical kinetics^10,16–18^. In our experiments the likely reasons for improved electron transfer kinetics can be attributed to the presence of nitrogen and oxygen functional groups that are deposited onto CNT surfaces by acid etching and electrochemical pretreatment. Nitrogen and oxygen have high affinities for hydrogen ions because of their net negative surface charges, hence the presence of these functional groups would have likely contributed to improved electrochemical kinetics. It has been shown that functional groups alter the electrochemical detection and the rate of electron transport on graphite surfaces, wherein groups having nitrogen and oxygen functional groups have superior electron transport kinetics^29^. Thus our work is consistent with previous observations.

Studies have shown that electrochemical oxidation of hydrogen peroxide on graphite surfaces occurs around 1.4V^28^. In our experiments we limited the waveform to 1.3 V. Despite our limitations, we were able to observe the deprotonation state of dopamine oxidation and the likely interactions between the intermediate species and water, which forms a peroxide-like species. These observations do conform to our hypothesis that using a hybrid electrode, i.e. CNT coating on pyrolytic-graphite with nitrogen doped graphite like electrode, we are able to show that 1) CNTs are able to offer faster electrochemical kinetics, wherein we successfully show the reversibility of dopamine reaction. 2) The ability of functionalized CNTs to store protons and offer reduced potentials. Our future work will focus on multimodal detection of dopamine and hydrogen peroxide using the “Dopamine Waveform”^25^. Limitations of our work is that we did not quantify the sensitivity of electrodes against scan frequencies. Nevertheless our modelling studies along with experimental work shows the heterogeneity of dopamine oxidation on CNT surfaces.

### 3. Selectivity and Specificity

The brain matter contains multiple analytes that tend to interfere with the determination of dopamine *in vivo*. These analytes compete for adsorption sites on the surface of the electrode. In addition materials can polymerise on the electrode over time, rendering it less sensitive. To address these issues chemical modification of electrodes was carried out with carbon nanotubes 5 mg mL^-1^ suspended in Nafion solution (dip-coating duration: 5 sec). The advantage of using Nafion as a selective membrane is the negative surface charge, therefore it allows selective passage of positive ions, dopamine, over negative ones, such as ascorbate^14,30^. To investigate the selectivity of pyrolytic-CNT coated electrodes, we introduced analytes which are routinely encountered in the brain during dopamine detection.

Using the “Dopamine Waveform”^25^, 1*μ*M solution of serotonin was introduced to the flow chamber, using T-400V s^-1^ at 10 Hz, waveform. Serotonin shows an oxidation peak at 0.22V and a secondary oxidation peak is seen around 1.0V (**figure 3A**). A reduction peak for serotonin complex at 0.4V, the kinetics of serotonin are not reversible, suggesting that the redox reactions for serotonin are far more complex than dopamine. We then investigated whether our electrodes and the scan parameters are suitable for serotonin detection, and would be able to withstand repetitive exposure to serotonin. (**Figure 3B**) shows the repetitive exposure of serotonin (10 cycles) onto the electrode surface. Multiple oxidation peaks are seen, wherein the oxidation peak of serotonin is shifted to left, suggesting the occurrence of biofouling of electrodes. A reduction peak can be seen at 1.0 V which is identical to that of dopamine detection, suggesting that it is from the reduction of peroxide like species which attacks indole groups, forming a compex, and polymerising it further. A reduction peak can be seen at 0V (**Figure 3B**), which is most likely an oxidised species being reduced to serotonin.

**Figure 3:**
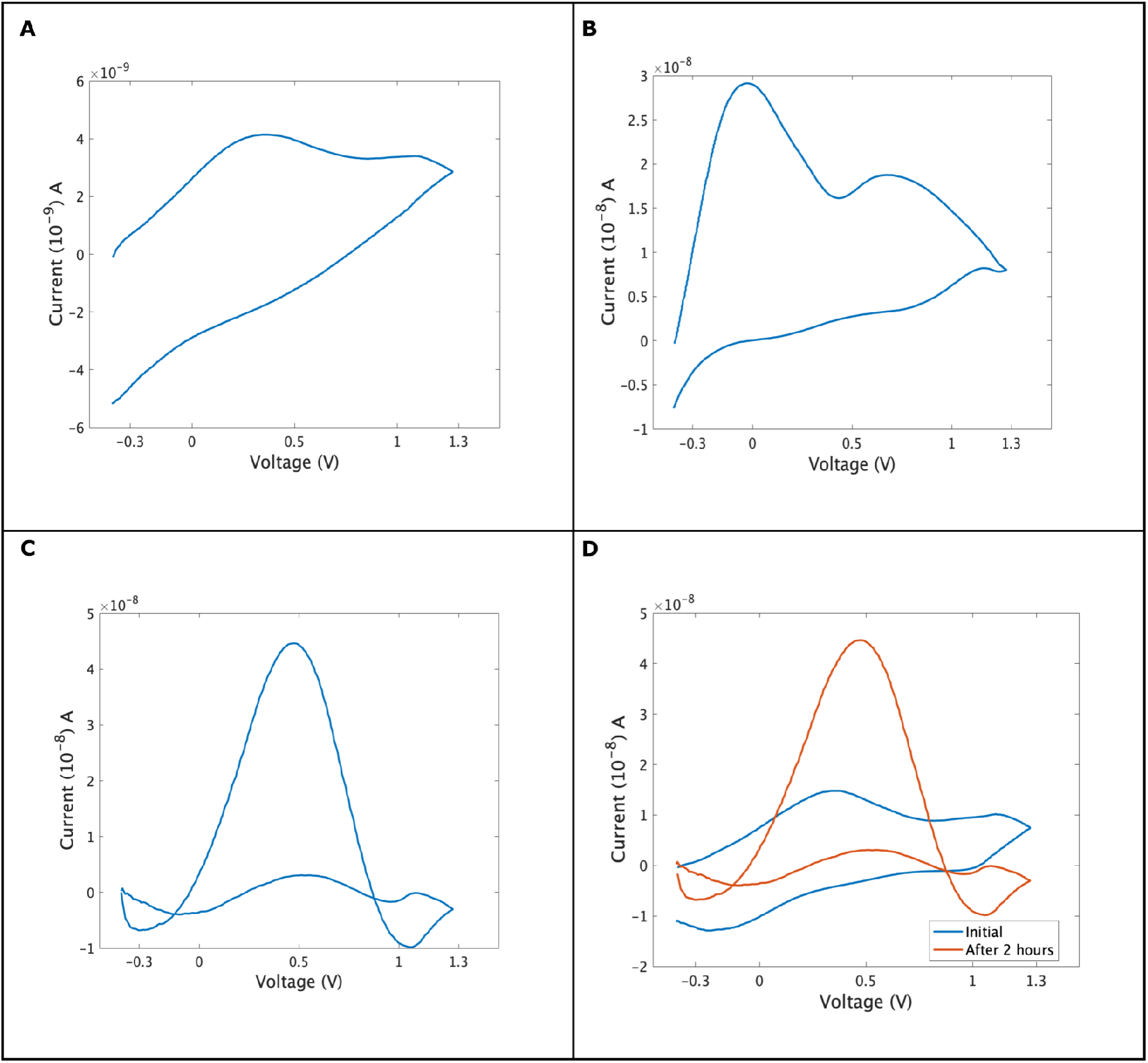
Cyclic voltammogram for analytes that are commonly occurring in the brain during dopamine detection. The waveform parameters were -0.4V to 1.3V, cycled back to -0.4V at 10 Hz at 400 V s^-1^. **A**: Shows cyclic voltammetry for serotonin 1*μ*M, showing just 1 oxidation peak at 0.29V and a broad reduction peak at 0.5V to 0.2V. **B**: Repeated exposure of serotonin to electrodes for 15 consecutive cycles. Two oxidation peaks can be seen for serotonin; where the peak at 0.75V is the oxidation peak and the at 0.2V is the reduction. An early oxidation peak can be seen as well, which can be an indication of biofouling of electrodes by serotonin polymerization on the surface of the electrode. **C**: Cyclic voltammogram of 3,4-Dihydroxyphenylacetic acid. Oxidation is seen at 0.45V and a broad reduction is seen at -0.1 V. **D**: Shows the oxidation of 1*μ*M dopamine at time point 0 and after 2 hours.

Studies that have made use of CNT coatings, suspended onto Nafion membrane and dip coated with graphite electrodes were able to detect serotonin without the occurrence of biofouling^22^. However they observed an overlap of oxidation peaks from dopamine and serotonin. The discrimintion of analytes was done based on the location of reduction peaks, wherein serotonin reduces around 0.1 V^22^. In our experiments we are able to discriminate between dopamine and serotonin based on the location of oxidation peaks and the shape and characteristics of voltagramm, however the occurrence of biofouling on our electrodes, limits our understanding of redox properties of serotonin. In the future we would make use of “Serotonin Waveform”^31–33^ for characterization of serotonin on CNT surfaces.

The byproduct of dopamine oxidation, 3,4-Dihydroxyphenylacetic acid (DOPAC) is present at a 100-1000 times higher concentrations than dopamine in the brain and has an oxidation potential identical to that of dopamine^14^. We cycled our electrodes in 10*μ*M of DOPAC and found that DOPAC oxidizes at 0.45V and reduces at 0.1V (**Figure 3C**). The electrochemical properties such as rise and fall time and oxidation potential of DOPAC is considerably different from those of dopamine, thus suggesting that both analytes can be differentiated using CNT electrodes. To understand if our probe is suited for *in vivo* neurochemistry and can withstand repeated exposure to dopamine we exposed our electrodes to 1*μ*M of dopamine for 2 hours (each exposure had 5 cycles). Initially, we found that dopamine was the predominant signal, however as time progressed we found a change in voltammogram characteristics thus suggesting that secondary analytes dominate the reaction (**Figure 3D**). Thus showing that our probes are more suitable for *in vitro* dopamine detection.

The small surface area of the electrodes provides more adsorption sites for analytes that are abundant in concentrations such as ascorbate, DOPAC and have higher diffusion coefficient (D). For our probes to selectively detect dopamine in the brain, the basal concentration at the site of detection must be higher than the surroundings, thus allowing dopamine molecules to diffuse at the surface of the electrodes, thereby causing Vroman effect^34–37^.

To selectively determine dopamine in presence of ascorbic acid, the electrode was exposed to a known concentration of dopamine and ascorbic acid (500*μ*M L^-1^). Oxidation peak current was noted for the analytes and the current was converted into a calibration factor (**Figure 4A**). As expected a linear increase in peak current occurs as concentration is increased (**Figure 4B**). The limit of chemical detection for dopamine is 50nM L^-1^, which is well below the basal physiological concentration of dopamine^22^. We then plotted the surface coverage of dopamine by exposing the electrodes to a range of concentrations. As the concentration of dopamine passes beyond 2*μ*M the linear behavior is lost although the peak current continues to increase with increasing concentration. To understand this the plot of peak current *vs* concentration, data was converted into a linearized version of Langmuir isotherms. At low concentrations the peak current is primarily due to adsorbed dopamine from bulk to surface of the electrode. As the concentration increases beyond 2*μ*M diffusion takes over so the current arises from a combination of preadsorbed dopamine on the surface and dissolved dopamine diffusing to the surface (**Figure 4C**).

**Figure 4:**
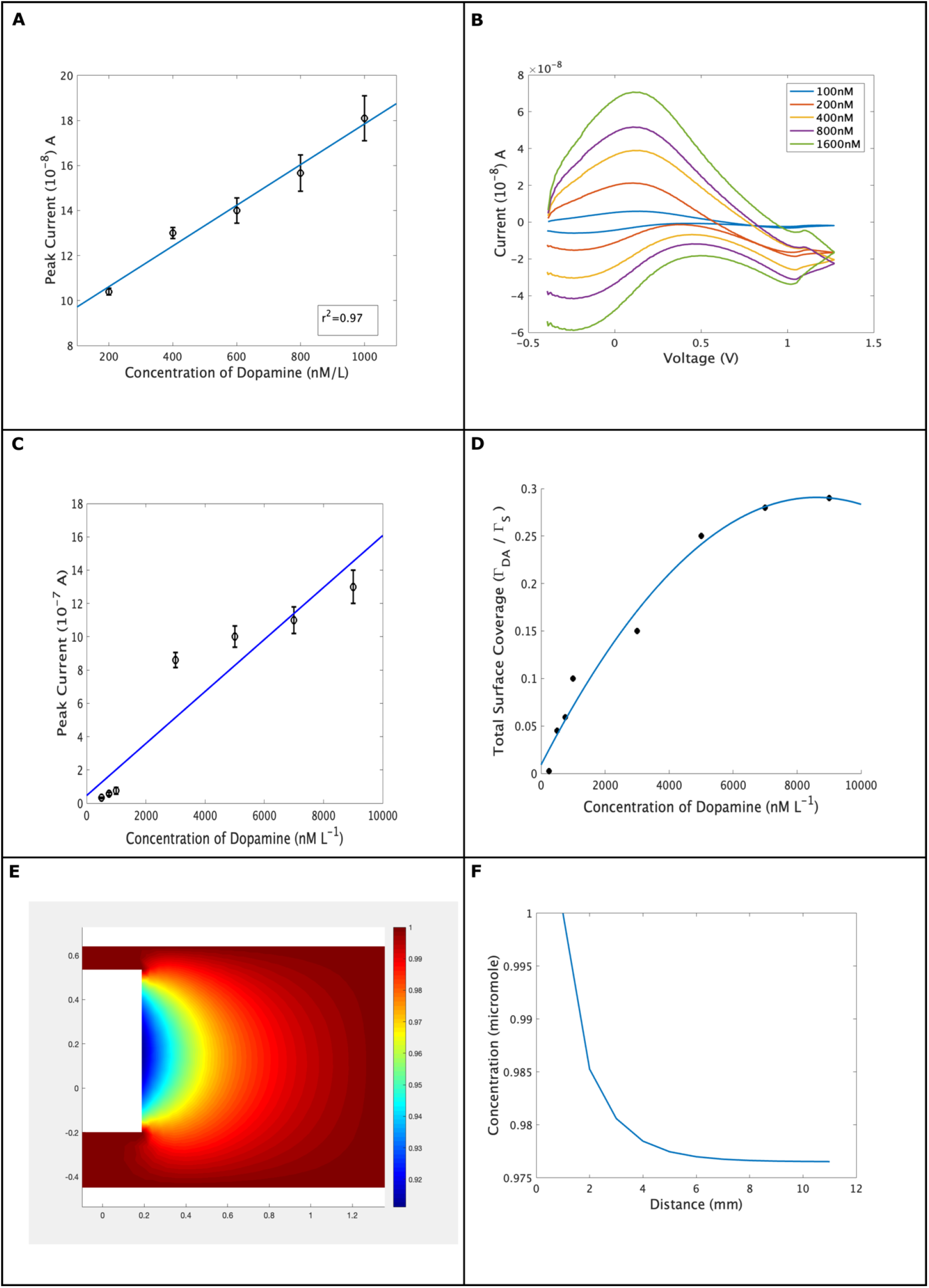
Shows the experimental and modeling response for dopamine on CNT electrodes. **A:** Shows a calibration curve for dopamine, where dopamine concentration varies from 50nM to 1600nM. The graph shows r^2^= 0.97. **B**: Shows overlaying of dopamine concentrations across trials. **C**: Shows the peak current for dopamine against a range of concentrations. **D**: The Langmuir-isothermal curve for dopamine against surface coverage^7,17,18,25^. **E**: Shows the modeling of dopamine from bulk to surface of the electrode. The concentration profile shows the strength of adsorption of dopamine on the surface of the electrode. **F:** The concentration profile as a function of distance (mm).

Even at higher concentrations the total surface area of the electrode is not saturated (**Figure 4D**) thus showing that the probes are well suited for studying a range of concentrations of dopamine covering those expected in diffusion-dominated processes, commonly occurring in the brain during volume transmission. To understand the mass transfer onto electrode surface from bulk to surface of the electrode, we modeled the edges of electrodes. The strength of adsorption is dependent on the surface area of the electrode, the position of the probe, and the diffusion coefficient of dopamine in Nafion (**Figure 4E and Figure 4F**). For 1*μ*M of dopamine in bulk, only a fraction is adsorbed on the surface of the electrode thus supporting our experimental hypothesis that total surface coverage might not be reached over small time constants, such as time course of neurotransmitter release, adsorption, and removal from the extracellular space.

### 4. Computational Model for Binding of Dopamine onto CNT Surface

To understand the interactions of dopamine with CNTs we model the interactions using ORCA^38^ workbench. We made use of Def2-SVP and def2-TZVP basis sets as they can accommodate larger atoms and provide convergence for bigger structures such as CNTs and the interactions of CNTs with dopamine^38,39^. CNTs optimization was done with CNTs capped with hydrogen and addition of oxygen as functional groups (**Supplementary Information I and II**). The interaction between dopamine and CNTs was given by means of dummy atoms (**Supplementary Information III**) as the iterations continue we saw that dopamine forms an electrostatic attraction with CNT in an aqueous medium, because of the dipolar nature of water (**Figure 5A**). The surface where CNTs were doped with oxygen shows the reduction of oxygen and liberation of protons (**Figure 5B and 5D**). The dipole interaction of water-CNT-dopamine prevents them from closing the bond distance and hence these molecules show a greater bond distance of (5.4 Å).

**Figure 5:**
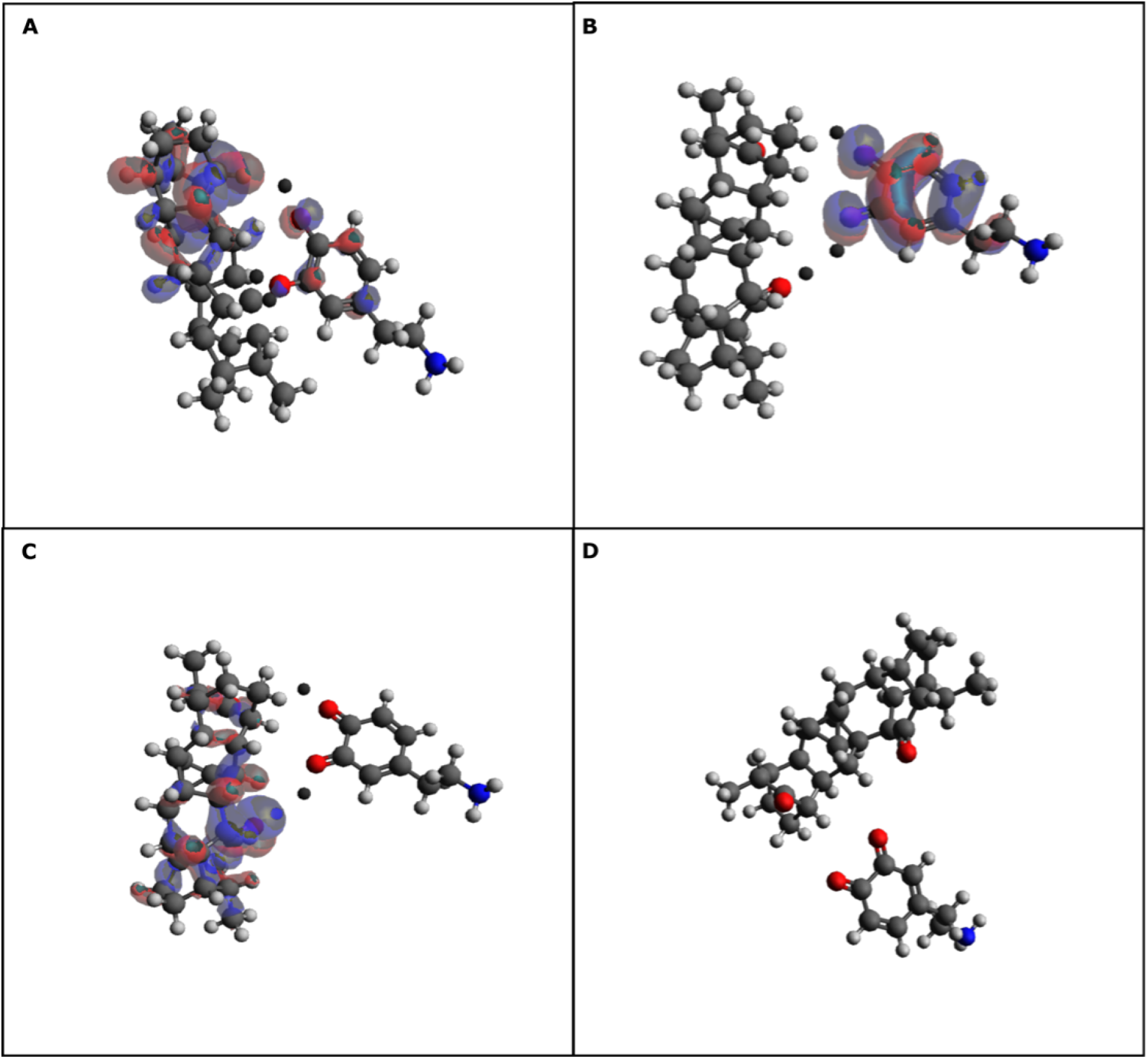
Shows the interaction of CNTs with dopamine. **A**: Dopamine interactions with CNT are limited when end groups are capped with hydrogen atoms. (Black spheres represent the dummy atoms). The bond distance between dopamine and CNTs is (2.8A). Dopamine oxidation occurs by proton and electron transfer. **B**: Shows the interaction of dopamine with oxygen along with HOMO and LUMO bond orbitals, wherein the intramolecular distance between them is (5.4 A). **C**: The interactions of CNTs with dopamine in LUMO(-1) orbitals. The oxidation of dopamine is seen with electron transfer at the hydroxyl group on dopamine. **D**: Shows the interaction of dopamine with CNTs capped with oxygen. The process is spontaneous and shows complete oxidation of dopamine.

In our present experiments we find that the interactions between CNTs and dopamine are limited and weak. Electrostatic bonds are weak in nature, thus the interaction between positively charged dopamine and negatively charged Nafion coated CNTs^30,40^ is a noncovalent one, which is consistent with our experimental observations. During our experimental work we found multiple oxidation states for dopamine (**Figure 1E**), thus confirming the likely hypothesis that oxidation of dopamine is a multistep process consisting of transfers of electrons and protons. The ability of CNTs to act as local proton storage, when doped with heteroatoms (**Figure 5C and 5D**) makes it possible to achieve reduced potentials for dopamine oxidation (**Figure 2D and Figure 2E**). Thus our experimental and modelling work shows that surface chemistry (**5A and 5B**) dictates the fate of electron transfer process which is consistent with experimental observations made on carbon fiber electrodes and glassy carbon electrodes by manipulating surface chemistry^29,41^.

DOPAC has a higher basal concentration in the brain compared to dopamine^14^. Given the higher basal concentration and chemical relatedness of DOPAC it is likely that DOPAC is predominantly detected over dopamine. DFT studies have shown that CNT electrodes have higher affinity for DOPAC than dopamine^42^. We tested this hypothesis by exposing our electrodes to known concentrations of DOPAC and found that DOPAC is oxidized at 0.5V and reduced at 0.1V (**Figure 3C**). Thus in our experimental work we find that DOPAC has a different oxidation potential as compared to dopamine (**Figure 1C and Figure 5D**), suggesting that heteroatoms play a significant role in modifying the potential. Indeed the peak amplitude is higher for DOPAC compared to dopamine, however this difference is attributed to the different concentration of analytes. Our future work will be developing the molecular model for adsorption of dopamine, DOPAC, serotonin and 5-hydroxyindoleacetic acid to understand the difference in conduction of molecules. In this way we are able to understand the kinetics of monoamines and their derivatives of oxidation onto CNTs electrodes.

The limitations of our computational model herein being the failure to construct a model in a unit cell with a well-defined slab. We did experiment with hexagonal, orthorhombic, and cubical geometry, however, our SCF convergence failed. We tried setting the maximum iterations to 500 in ORCA, however, this did not improve our convergence. Hence we adopted convergence in the open-space model, wherein the solvation properties, current, and temperature were defined globally. We did attempt to show the binding of dopamine to the basal plane of CNT, however, this convergence failed (data not shown) as well, suggesting that the binding of dopamine-CNT surface can occur only laterally and not on basal planes. While we did not take into account the CNT density, i.e. binding of one molecule of dopamine to multiple CNT molecules..

## Conclusions

In this work, we show that doping of pyrolytic carbon electrodes with functionalized CNTs improves the electrochemical kinetics for dopamine. We were able to perform multimodal detection of dopamine and hydrogen peroxide like species, formed during oxidation of dopamine in the aqueous solution. These CNT coated electrodes are able to detect dopamine at higher sampling frequency by improving the rate of electron transfer, thus allowing us to show the reversible nature of dopamine electrochemical reaction. The ability of CNT coated electrodes to detect nanomolar concentrations of dopamine in the brain at sub-millisecond timescale makes them a suitable candidate for studying neurochemical communication in the brain. Our future work will be detecting dopamine in the biological medium.

## Materials and Methods

All chemicals, dopamine hydrochloride (DA-HCl), serotonin hydrochloride (5-HT-HCl) ascorbic acid, 4,-(2-hydroxyethyl-)1-piperazineethanesulfonic acid (HEPES), single-walled carbon nanotubes (SWCNTs), sodium chloride (NaCl), potassium chloride (KCl), sodium bicarbonate (NaHCO_3_), magnesium chloride (MgCl_2_), monosodium phosphate (NaH_2_PO_4_), Aluminum oxide (Al_2_O_3_)(10μm) and 3,4-Dihydroxyphenylacetic acid (DOPA), were purchased from Sigma Aldrich (Germany) and were of analytical grade. Quartz capillaries (o.d. 1 mm, i.d. 0.5mm, length 7.5 cm were purchased from Sutter Instruments, Germany.

### 1. Electrochemical setup

Electrochemical detection of dopamine was done using a two-electrode system that was connected to a patch-clamp amplifier (Intan Technologies, Los Angeles, USA). The working electrode consisted of graphite electrodes coated with CNTs and the reference electrode was silver wire coated in KCl solution (3.5M). The assembly was placed in a Faraday cage to shield it from external noise. Electrochemical recordings were done in a physiological chamber that was able to mimic the transient change in neurotransmitter concentrations^43,44^. Solutions were delivered to the recording chamber at 32°C to mimic a brain-like environment. Buffer was delivered at a flow rate of 2mL min^-1^ and analyte was introduced as a bolus on 5 s, followed by 10 s pause^10^. A voltage waveform was applied from -0.4V to +1.3V at a scan rate of 400V s^-1^ at 10Hz, with a step size of 5mV. To obtain a stable background current, electrodes were scanned in a buffer for 30-45 min. at a scan rate of 1000V s^-1^. For electrochemical measurements of neurotransmitters, quartz capillaries were heated for 60 s, dipped in CNTs dispersed in Nafion solution (5mg mL^-1^ in ethanol). To obtain the Faradaic contributions from dopamine, the background was subtracted^45,46^ digitally using a custom MATLAB script. Electrochemical measurement of dopamine was carried out in artificial cerebrospinal fluid (aCSF) as described before^47–50^ and included 135mM NaCl, 5.4mM KCl, 5mM Na-HEPES buffer, 1.8mM CaCl_2_ and 1mM of MgCl_2_.

### 2. Scanning Electron Microscopy of Electrodes

Scanning electron microscopy was performed using a JEOL 6330 Cryo FESEM. Samples were cut into 2cm lengths and coated with a gold layer to prevent charging.

### 3. Preparation of ultramicroelectrodes

To obtain micrometer quartz capillaries, a CO_2_ laser puller was used (Sutter instruments). The laser beam produced heat of 750°C, using the pulling velocity of 50 and a delay (DEL) of 127, to give a final tip diameter of 0.8-1.5μm. The tip was manually shortened to reach a final diameter of 25-30μm.

### 4. Pyrolysis

Carbonized microelectrodes were fabricated by performing pyrolysis of propane in a nitrogen environment in a quartz microcapillary^51,52^). The heat was delivered at the narrower end using a propane-butane torch for 60 seconds. To improve the electric contact carbon was deposited on the broader end of the quartz capillary. After the pyrolysis was done, pyrolytic carbon capillaries were allowed to cool for 2 min under a nitrogen environment. Following the pyrolysis, electrodes were coated with CNT solution suspended in Nafion.

### 5. Computational Models

Computer models for CNTs and dopamine were constructed in Avogadro^53^ supported with ORCA^38,54^. CNTs model consisted of a small chain of CNTs (2,1) where the ends and sides were capped with hydrogens as an example. Geometry optimization of CNTs was done using DFT theory calculated in ORCA^38,54^ using def2-SVP and def2-TZYVP basis sets^39^(**Supplementary Information I and Supplementary Information II**). To understand the binding of dopamine onto the CNTs surface, first dopamine’s geometry was optimized using implicit solvent mode in ORCA. Then dopamine-CNTs interactions were visualized To facilitate the transition search, dopamine and CNTs were connected through dummy atoms and no bond or distance constraints were laid (**Supplementary Information III**). The structures were optimized in ORCA and the resultant output file was loaded in Avogadro for visualization.

### 6. Theory

To model the adsorption of dopamine onto the surface of the electrode, a finite-difference simulation was carried out using the ODE toolbox from MATLAB (MathWorks, USA). The system consisted of a 2-D model wherein the edges/planes of the electrodes were modeled. This approach allowed us to address the mass deposition on the surface, which was governed by the diffusion of dopamine from bulk to the surface of the electrode via adsorption^25^. Fick’s second law of diffusion^55^ was expressed in a time-resolved manner, where the relationship between the diffusion coefficient(D), concentration (C), and the Laplacian operator is expressed as

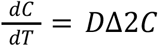

In our experiments, we assume that the diffusion constants for reactants, products, and intermediates are identical. The Laplace operator is given by the geometry of the electrode^56^. The current generated (*I*) at the surface of the electrode by application of a potential is,

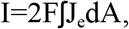

Where J_e_ is the flux at the electrode surface caused by adsorption and desorption of dopamine^25^. The relations between dopamine adsorption energy on the surface of the electrode and the surface coverage can be given by the Langmuir equation^18^, wherein for a low concentration of dopamine the plot is linear. As the concentration increases the adsorption sites are not increased thus requiring a linearized version of the Langmuir equation, which can be given by surface coverage

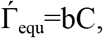

Where b is the strength of adsorption and the flux at the surface of the electrode can be given by rapid adsorption and desorption of dopamine on the surface of the electrode^18^

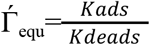

K_ads_ and K_deads_ are the kinetic rate constants that are determined experimentally^56^.

**Figure 1:**
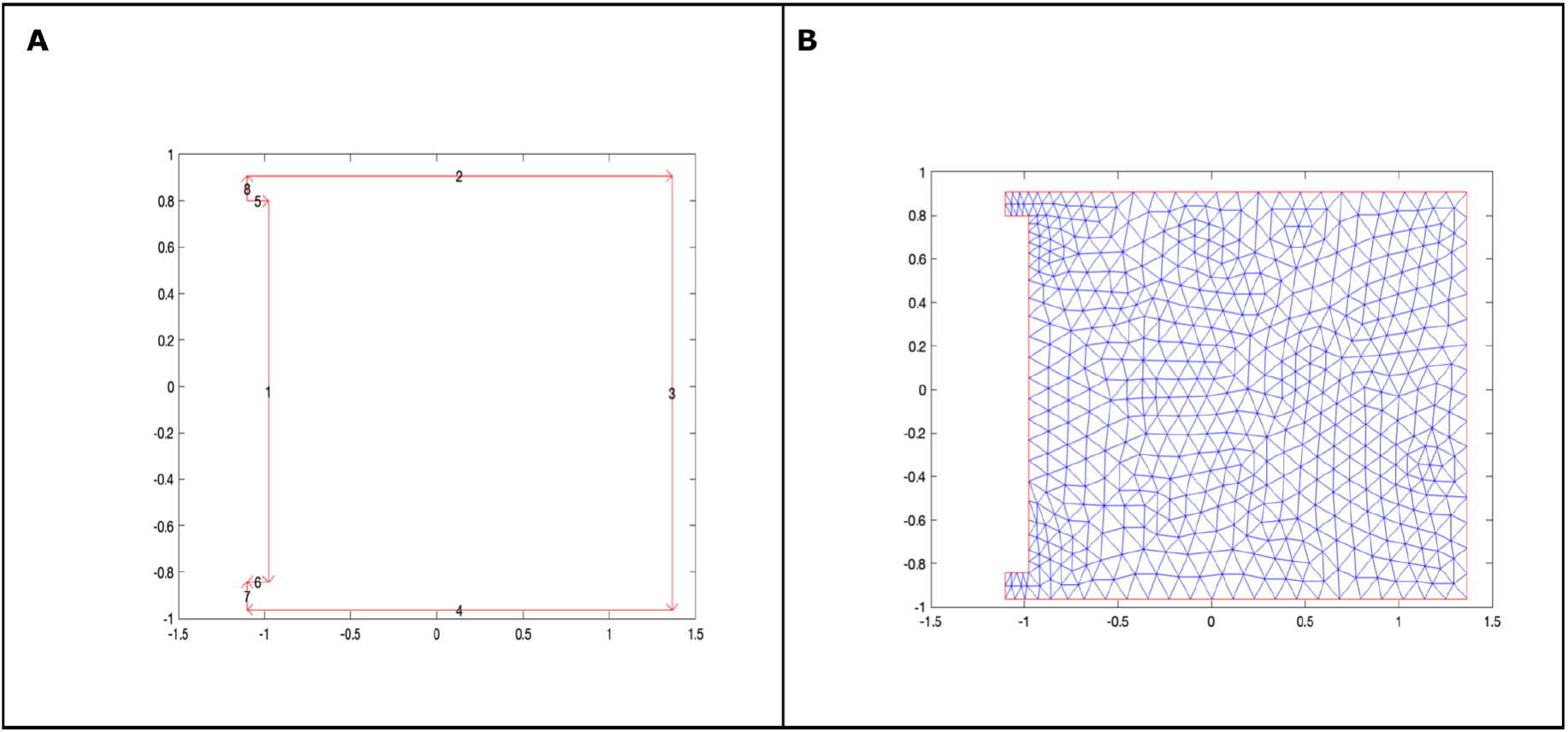
A 2-D view of electrode geometry in the Cartesian Coordinates. **A**: The electrode is modeled in multiple edges (1--8) as shown. Diffusion is only occurring on face 1, which is described by Neumann boundary conditions while all other boundaries are in Dirichlet boundaries. **B:** To facilitate the modeling of the diffusion problem, the electrode face is finely meshed (**Supplementary Figure-1**) and diffusion is only described on the mesh points that are only the edges of plane 1.

## Acknowledgements

We would like to thank Prof. Frank Platte for his help with ODE modelling. We would also like to thank Geert-Jan Janssen for his help with electron microscopy. This work was supported in part by European Regional Development Fund (MIND, nr. 122035).

## Supplementary Information

**Supplementary figure 1.**
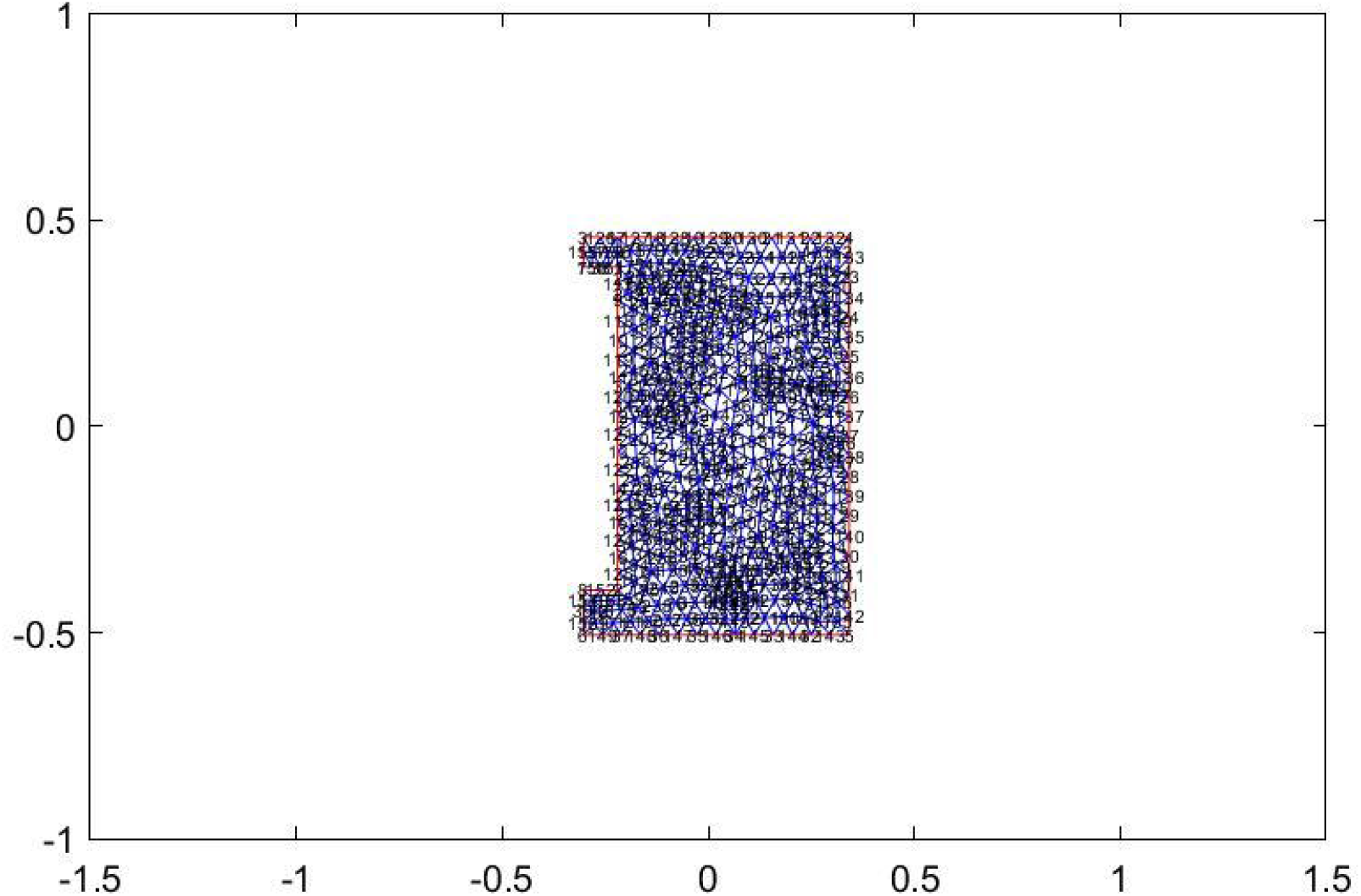
A 2-D model showing the edges of the electrodes with mesh numbers. Diffusion is described only from the points (117-126).

## Supplementary Information-I

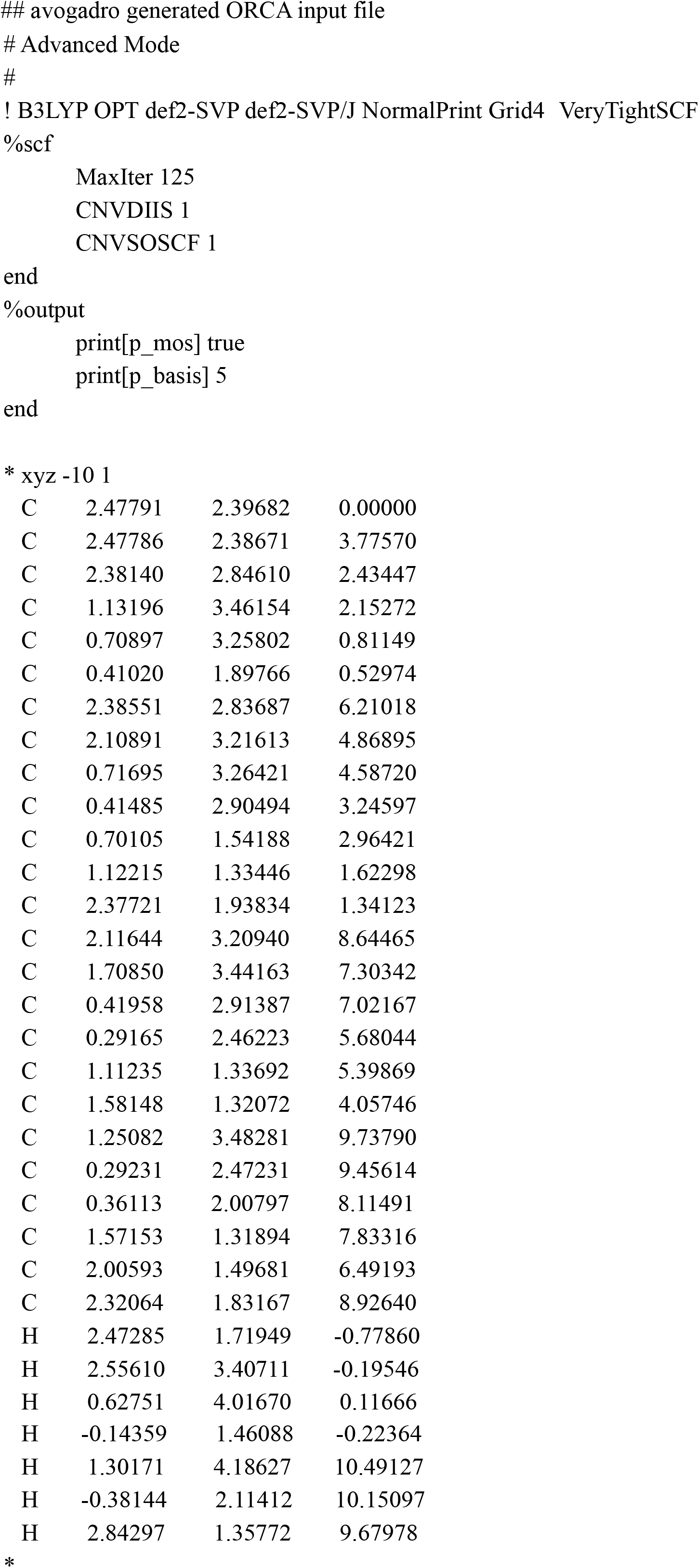

## Supplementary Information-II

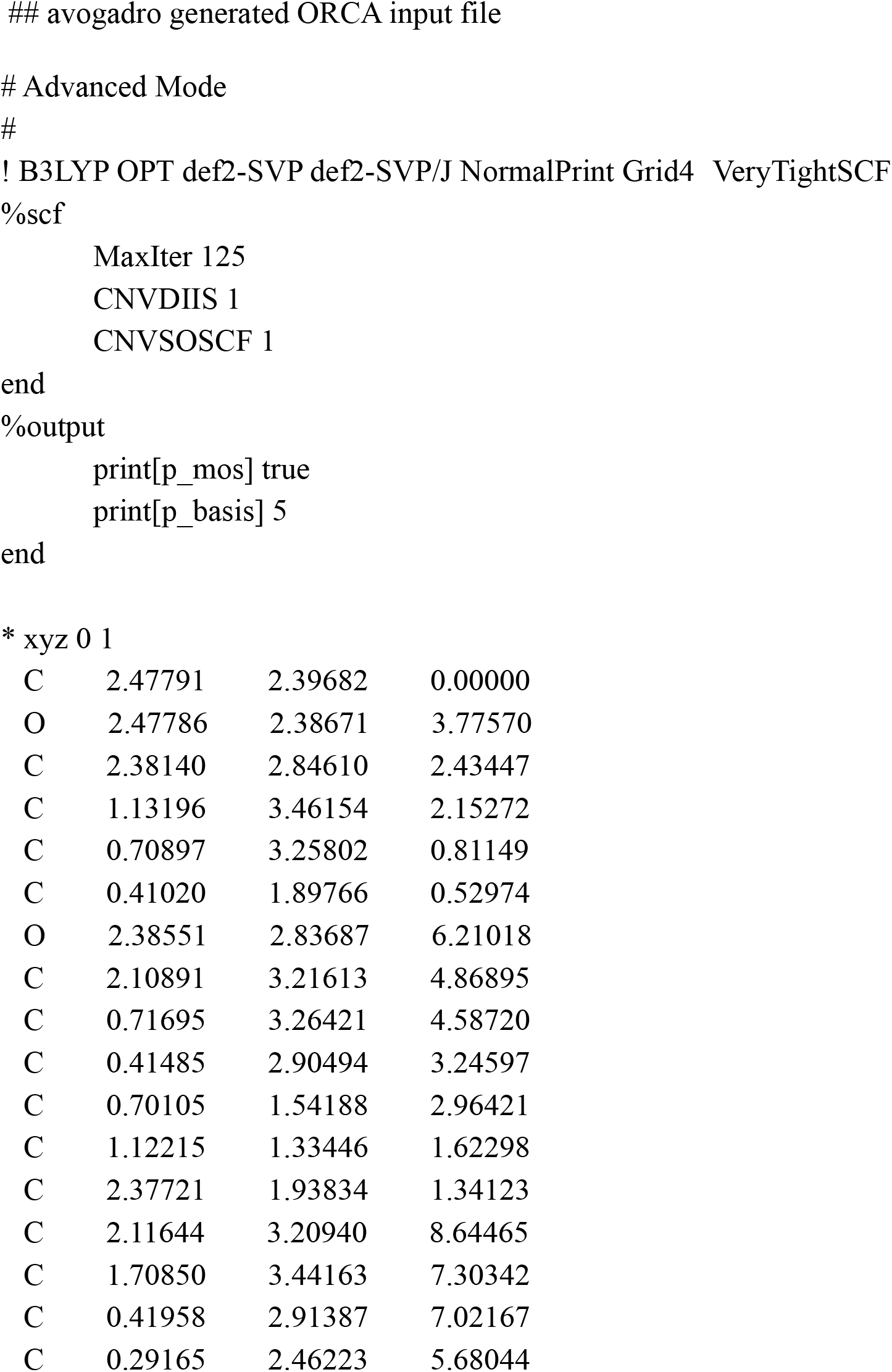
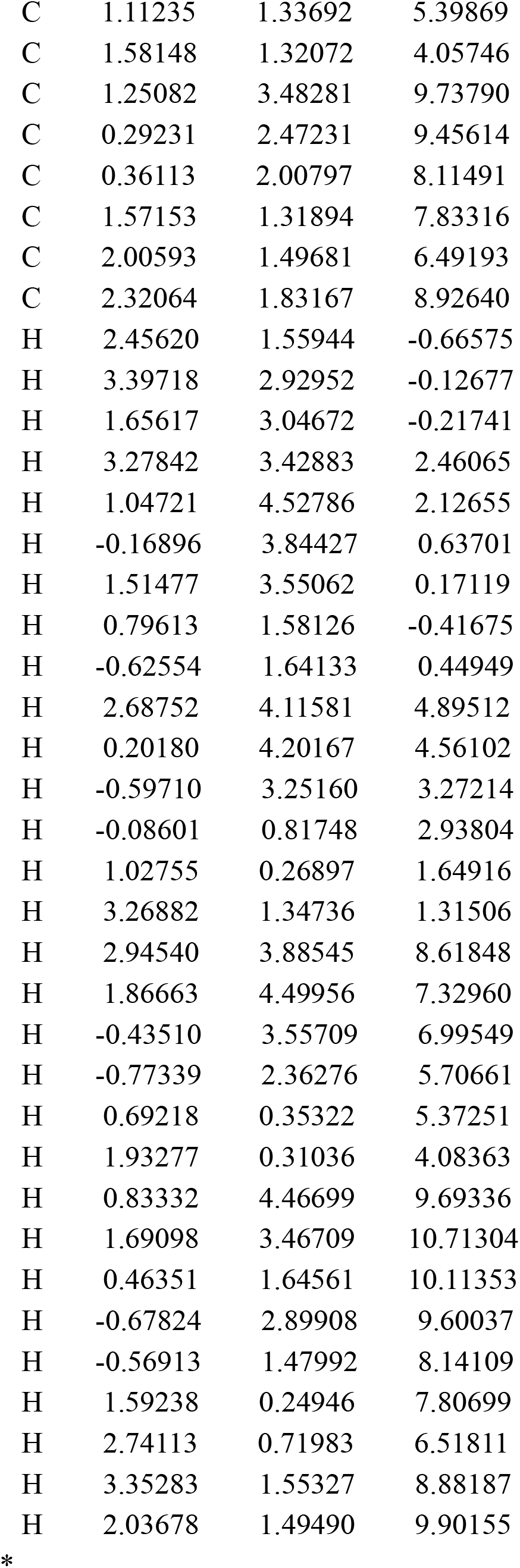

## Supplementary Information III

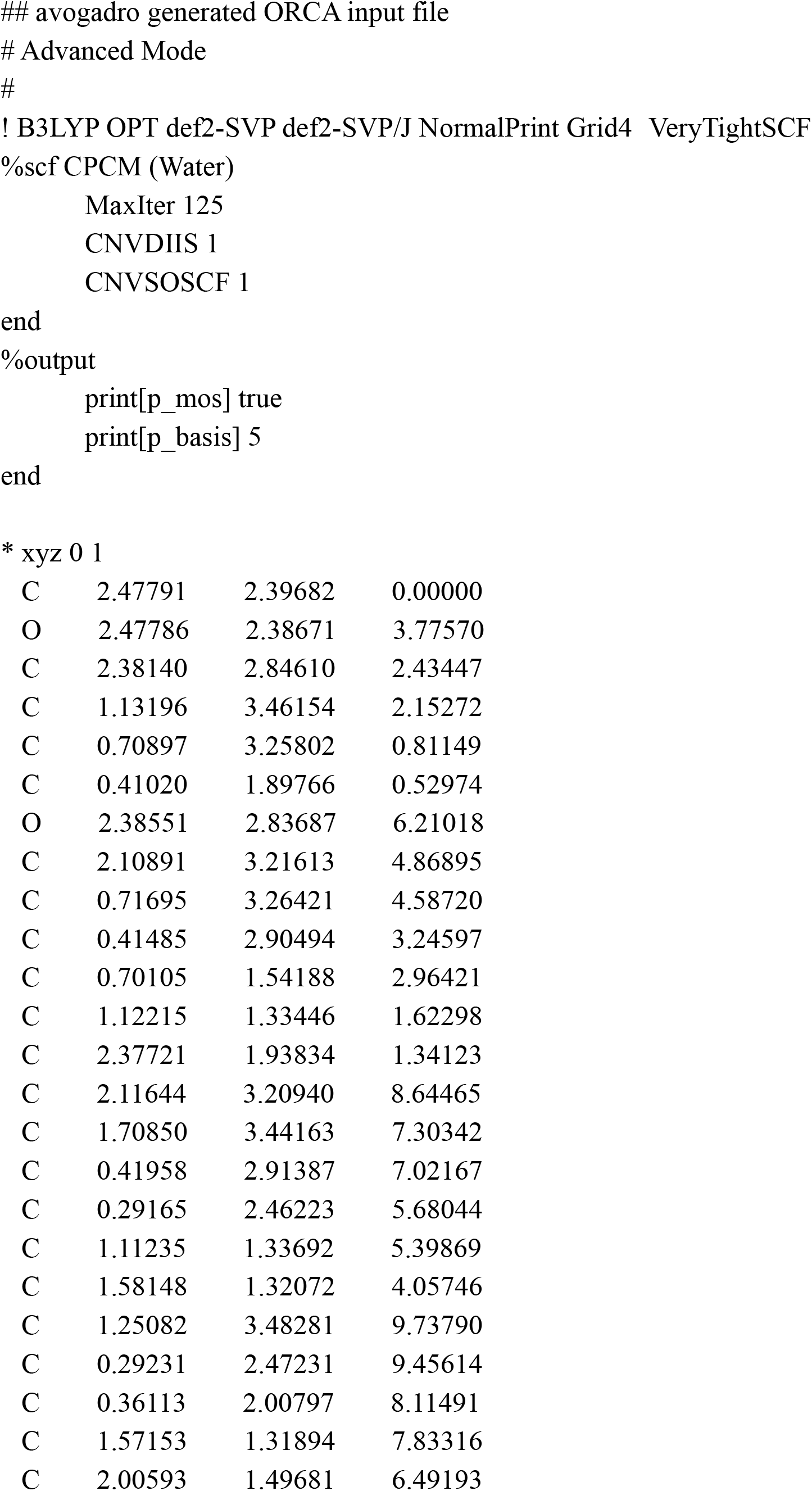
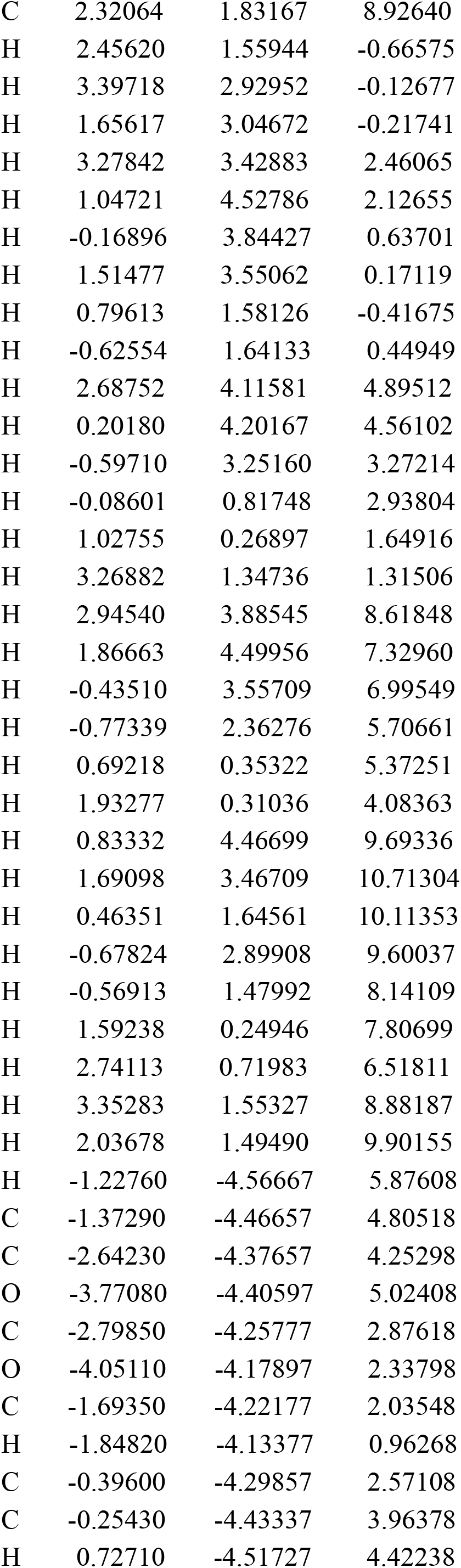
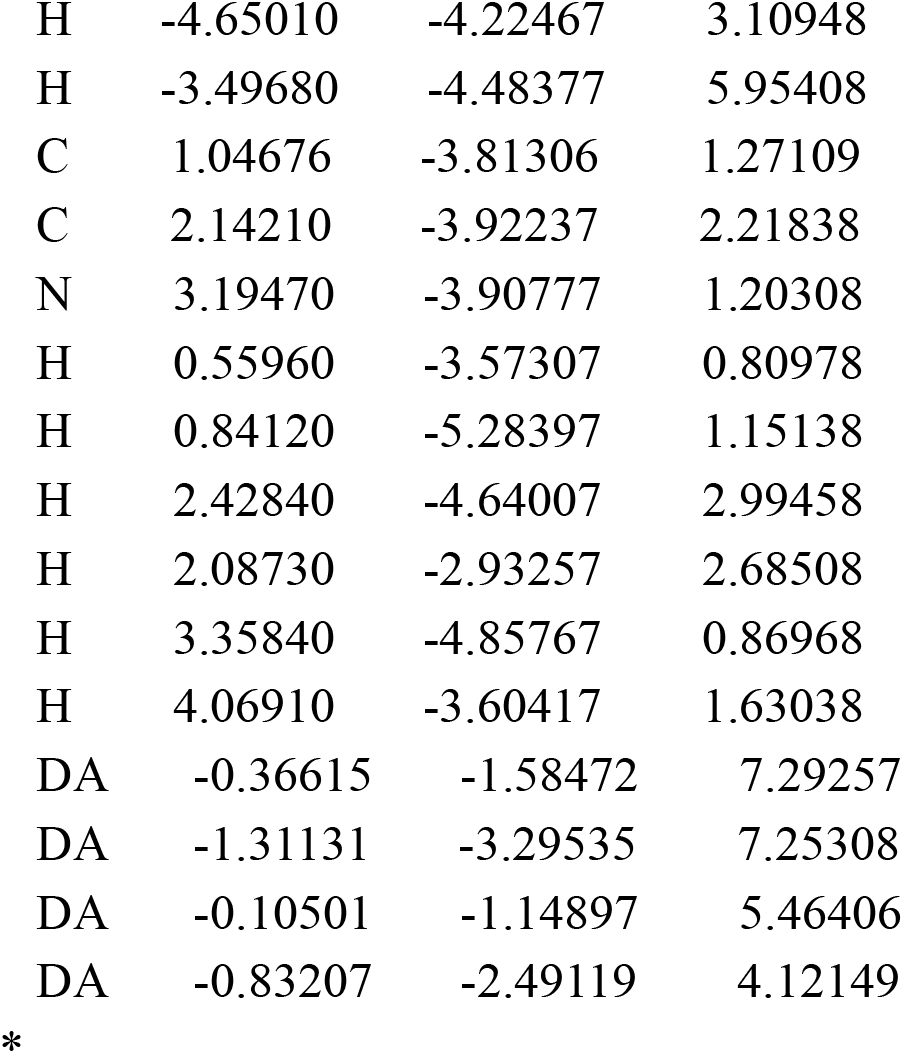

